# Effects of lipid composition on membrane distribution and permeability of natural quinones

**DOI:** 10.1101/568816

**Authors:** Murilo Hoias Teixeira, Guilherme Menegon Arantes

**Affiliations:** Department of Biochemistry, Instituto de Química, Universidade de São Paulo, Av. Prof. Lineu Prestes 748, 05508-900, São Paulo, SP, Brazil

## Abstract

Natural quinones are amphiphilic molecules that function as mobile charge carriers in biological energy transduction. Their distribution and permeation across membranes are important for binding to enzymatic complexes and for proton translocation. Here, we employ molecular dynamics simulations and free energy calculations with a carefully calibrated classical force-field to probe quinone distribution and permeation in a multicomponent bilayer trying to mimic the composition of membranes involved in bioenergetic processes. Ubiquinone, ubiquinol, plastoquinone and menaquinone molecules with short and long isoprenoid tails are simulated. We find that water penetration increases considerably in the less ordered and porous bilayer formed by di-linoleoyl (18:2) phospholipids, resulting in a lower free energy barrier for quinone permeation and faster transversal diffusion. In equilibrium, quinone and quinol heads localize preferentially near lipid glycerol groups, but do not perform specific contacts with lipid polar heads. Quinone distribution is not altered significantly by the quinone head, tail and lipid composition in comparison to a single-component bilayer. This study highlights the role of acyl chain unsaturation for molecular permeation and transversal diffusion across biological membranes.

## Introduction

Electron transfer chains (ETC) involved in biological energy transduction rely on natural quinones and their redox-coupled quinols (Q molecules^†^, Figure 1) to shuttle electrons between protein complexes and translocate protons across phospholipid membranes, contributing to generation of an electrochemical gradient.^1–4^ Clearly, the mechanisms for molecular recognition and binding of Q by respiratory and photosynthetic enzymatic (super)complexes^5,6^ depend on the membrane distribution of Q molecules.^7–9^

**Figure 1:**
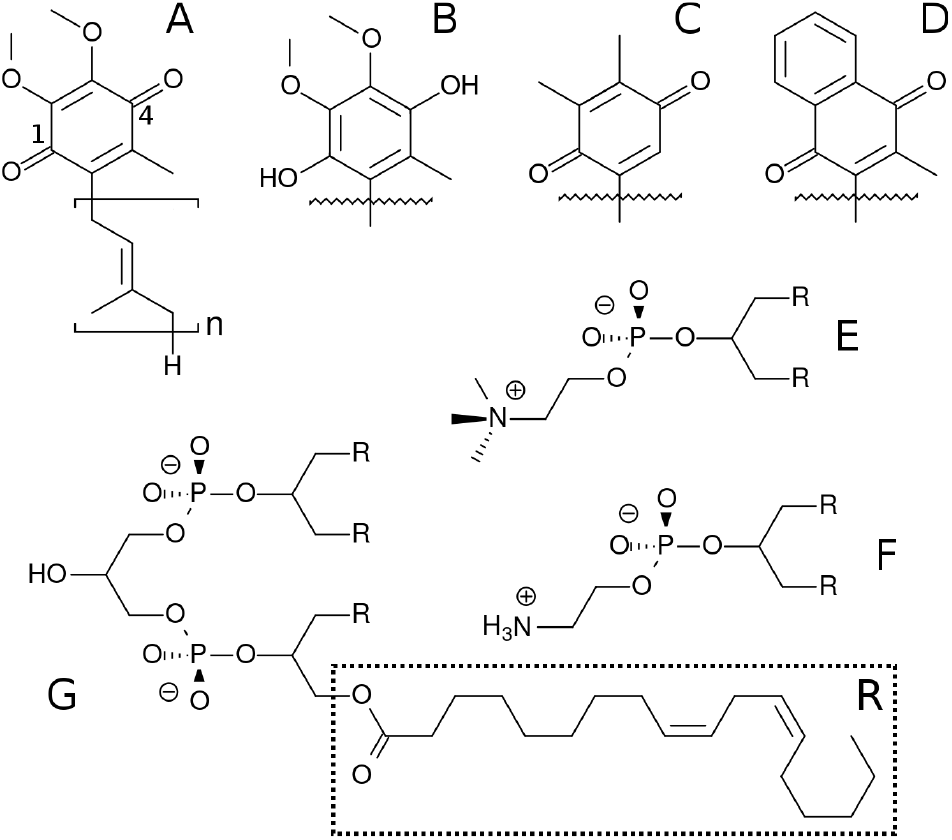
Chemical structure of Q molecules and lipids studied here. A is ubiquinone (UQ_*n*_), B is ubiquinol (UQ_*n*_H_2_), C is plastoquinone (PQ_*n*_), D is menaquinone (MQ_*n*_). The n subscript is the number of bound isoprenoid units, shown only for UQ_*n*_ which also displays positions 1 and 4 of the Q-ring. E is 1,2-dilinoleoyl-sn-glycero-3-phosphatidylcholine (DLPC), where R is the linoleoyl (18:2) acyl chain. F is 1,2-dilinoleoyl-sn-glycero-3-phosphatidylethanolamine (DLPE) and G is 1 ‘-3’-bis[1,2-dilinoleoyl-sn-glycero-3-phospho]-sn-glycerol (cardiolipin, LCL).

Diffusion of Q on the membrane has also been suggested to control ETC turnover rates.^10–13^ In redox loops translocating protons across the membrane directly through a quinone/quinol pair (or pool), as in the Q-cycle,^14,15^ at least two events of Q permeation, or transversal diffusion, occur. In the mitochondrial ETC, quinol production in respiratory complexes I, II and III take place at the membrane N-side, but the Q_*o*_ site for quinol oxidation in complex III is at the P-side.^16–18^ Alternative oxidases cytochrome *bo*^19^ and cytochrome *bd*^20^ in bacteria also carry out quinol oxidation at the membrane P-side. Thus, quinone/quinol recycling depends on Q permeating the membrane back-and-forth and it is important to understand this molecular process in detail.

Most of these proposals focused on Q lateral diffusion over the membrane, a fast and essentially diffusional process.^21^ But transversal diffusion is an activated process and hence much slower.^22–25^ Unfavorable desolvation and dielectric interactions (with associated energetic barriers) have to be overcome when the polar Q-head penetrates the lipid bilayer.^18,26^

Membrane distribution and lateral diffusion of Q have been experimentally studied (and debated) for decades.^21,25,27–32^ Transversal diffusion is harder to measure accurately and has received less attention.^24,25^ Molecular dynamics (MD) simulation is a powerful technique to probe both membrane distribution and permeation, but the quality of the results heavily depend on the force-field description of molecular interactions.^33,34^ Recently, MD simulations were used to investigate the permeation of ubiquinone on a 1-palmitoyl-2-oleoyl-sn-glycero-3-phosphatidylcholine (POPC) single-component bilayer and found the Q-head preferentially localizes near lipid glycerol groups.^26^

Lipid composition modulates the properties of membranes and embedded molecules.^35,36^ For instance, the fraction of unsaturation (double bond) in lipid acyl chains changes membrane fluidity and permeability so that cells have evolved to homeostatically control this property through the biosynthesis of fatty acids.^36,37^ Bioenergetic membranes are enriched with the anionic cardiolipin and its role on the distribution of ubiquinone in a more realistic multi-component lipid bilayer has also been explored by simulation.^38^

Here, we employ MD simulations with a force-field carefully calibrated for Q molecules^26^ to investigate the effects of lipid composition and unsaturation of acyl chains on Q distribution and permeation using a multi-component bilayer mimicking membranes involved in bioenergetic processes. Ubiquinone, ubiquinol, menaquinone and plastoquinone (Fig. 1) are studied to probe the role of the Q-head on membrane localization.

## Methods

### Set-up for molecular dynamics simulations

Six model bilayers were studied here, composed by 240 DLPC (Fig. 1), 208 DLPE, 64 cardiolipins (LCL di-anion), 16 Q (UQ_2_, UQ_2_H_2_, UQ_10_, UQ_10_H_2_, PQ_9_ or MQ_9_) molecules symmetrically divided between each leaflet, 128 Na^+^ cations and 21728 water molecules. This is the same lipid composition used previously by Róg *et al*.^38^ In umbrella sampling simulations, 37435 water molecules were used.

The initial configuration for the UQ_10_H_2_ bilayer was kindly provided by Prof. Tomasz Róg as used in their publication.^38^ Initial geometries for all other bilayers were derived from this configuration by deleting or “mutating” Q-head atoms and adapting the atomic connectivity accordingly.

Force-field parameters for lipids and Na^+^ were taken from CHARMM36.^39^ Water was described by TIP3P.^40^ Our CHARMM^41,42^ compatible force-field previously obtained by careful calibration^26^ was used for Q molecules. Additional parameters necessary for MQ and PQ heads were taken directly from the CHARMM specification with partial charges and atom types shown in Table S1. Complete topologies and parameters are available from the authors upon request.

All MD simulations were carried out with GROMACS^43^ versions 4.6.7 and 5.1.3. The NPT ensemble was used and temperature kept at 310K with the Bussi thermostat^44^ and a coupling constant of 0.1 ps with two separate coupling groups (water and everything else). Pressure was kept at 1.0 bar with a weak coupling scheme with a compressibility of 0.5-1.0 10^−5^ bar^−1^ and a coupling constant of 1 ps. Semi-isotropic coupling in the direction normal to the bilayer was applied. Electrostatics were handled by PME method^45^ with a real space cutoff of 1.2 nm, grid spacing of 0.13 nm and quartic interpolation. All covalent hydrogen bonds were constrained using LINCS^46^ and van der Waals interactions were truncated from 1.0 to 1.2 nm. No dispersion corrections were applied.^47^ The integration time step was set to 2 fs and the nonbonded list was updated every 20 fs.

Unrestrained MD simulations were performed for 300 ns, frames were recorded every 20 ps and the initial 50 ns were discarded to allow for equilibration in the trajectories analyzed here. Mean area was computed as the ratio between the average area of the membrane plane and the number of lipid heavy atoms per leaflet. Atoms instead of molecules were used to allow comparison with multi-component bilayers. Normalized mass densities were calculated to ease comparison between groups with different number of atoms. Contacts between Q atoms and solvent molecules were defined with a 0.3 nm cut-off. Order parameters for methylene units of acyl chains were also calculated.^48^

### Free energy calculations

Umbrella sampling (US)^49^ was used to compute the free energy profile for UQ_2_ permeation across the membrane normal. The distance between the UQ_2_ head center-of-mass (COM) and the membrane COM along its normal (*z*-axis) was used as the reaction coordinate. The membrane COM was computed from a sum over the lipid atoms within a cylinder centered in the quinone and a 2.0 nm radius. This cylinder scheme helps to avoid artifacts due to membrane undulations, which are common in large membranes.^50^

Each simulation box contained 16 UQ_2_ molecules, so 16 different windows were computed simultaneously in each US simulation. To avoid spurious interactions between UQ_2_ molecules (for instance, aggregation when in the water phase), an artificial repulsive interaction was included between each Q-head C6 atom pair, using a specific Lennard-Jones pair with zeroed dispersion. Initial configurations for US were obtained at *t* ~ 200 ns from the UQ_2_ unrestrained MD trajectory described above.

US windows were chosen equally spaced by 0.250 nm in the ranges *z*=[0.125,1.375] nm and [3.10,4.35] nm, and spaced by 0.125 nm in the range *z*=[1.500,2.875] nm. The umbrella potential was set with *k_umb_*=1000 kJ mol^−1^ nm^−1^ and a total of 48 windows (obtained from 3 separate US simulations) were used to cover UQ_2_ permeation over the complete membrane normal (24 windows for each leaflet). Each US simulation was ran for 300 ns. The reaction coordinate was recorded every 0.2 ps. Free energy profile was symmetrized and calculated with WHAM.^51,52^ The initial 30 ns were discarded for equilibration and the remaining 270 ns were used for data accumulation on each window. Statistical uncertainties were estimated as 95% confidence intervals by bootstrap analysis.^53^

## Results & Discussion

### Quinone composition and multi-component lipid membranes do not change the Q-head equilibrium distribution

Results from unrestrained molecular dynamics simulations for six membranes with different Q composition are presented in this section. Mass density profiles for lipid groups are shown in Fig. 2. Density distributions in each leaflet are fairly symmetrical (negative or positive membrane normal) which indicates good equilibration and sampling of MD trajectories. Densities of phosphate (P) and acyl glycerol (shown only in Fig. 2A) groups are unaltered in all membranes and suggest that Q composition does not perturb the membrane structure.

**Figure 2:**
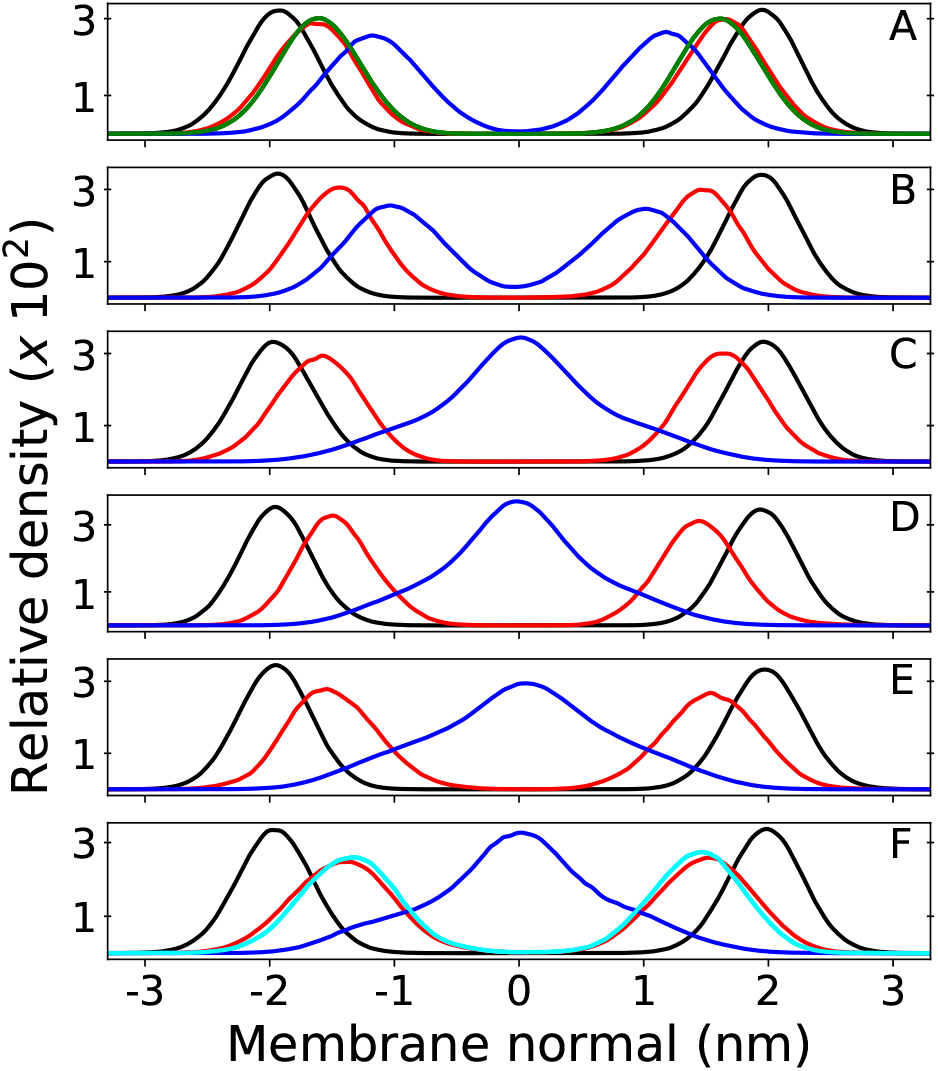
Relative mass density of multicomponent bilayers with different quinone analogue (Q) composition. Phosphate group of DLPC+DLPE is shown in black, Q-head in red and Q-tail in blue. Panel A shows the bilayer containing ubiquinone-2 (UQ_2_), where the green profile indicates density of the DLPC+DLPE glycerol group; B is ubiquinol-2 (UQ_2_H_2_); C is ubiquinone-10 (UQ_10_); D is ubiquinol-10 (UQ_10_H_2_); E is plastoquinone-9 (PQ_9_) and F is menaquinone-9 (MQ_9_), where the cyan profile indicates density of the first Q-ring (6 C atoms) only and the full Q-head (10 C atoms) is in red. Zero of the membrane normal corresponds to the bilayer center. Data were not symmetrized between the two leaflets.

Density peaks for the Q-head change only 0.3 nm (from 1.4 to 1.7 nm in modulus) between all Q. Density peaks indicate the Q-head equilibrium position across the membrane normal and are in agreement with equilibrium positions previously determined for ubiquinone and ubiquinol in POPC membranes.^26^ Q-head density distributions also do not depend on isoprenoid chain length (for instance, compare Fig. 2A and C). For short isoprenoid chains (*n* = 2), the Q-tail density localizes around 1 nm. For longer chains (*n* = 9, 10), the Q-tail density peaks at the membrane center and spreads through most of the lipid phase. UQ_*n*_H_2_ (*n* = 2, 10) is slightly more buried, with Q-head peak density 0.2 nm lower than ubiquinone and with more penetration of Q-tail density towards the membrane center. This behavior was also observed in POPC membranes and is a consequence of the intramolecular hydrogen bonds (H-bond) often observed between phenolic hydrogen and methoxy oxygen in ubiquinol.^26,54^ These H-bonds stabilize the Q-head when it is desolvated and allow for a higher permeation of UQ*n*H_2_.

Fig. 3 shows contacts performed by the Q-head O1 with different lipid groups and solvent. H-bond acceptors display shorter water contacts as the minimum distance shown in Fig. 3E is computed between O1 and any of the water atoms. More interestingly, MQ_9_ shows the smallest first hydration peak and only UQ_10_ and MQ_9_ show a secondary water peak at a distance of ~0.4 nm, corresponding to configurations that O1 is not forming an H-bond with water but the other ketonic O4 is. Similar distributions but with smaller differences between Q molecules are observed for contacts with the Q-head O4, except for MQ_9_ which is significantly more hydrated in O4 (data not shown here).

**Figure 3:**
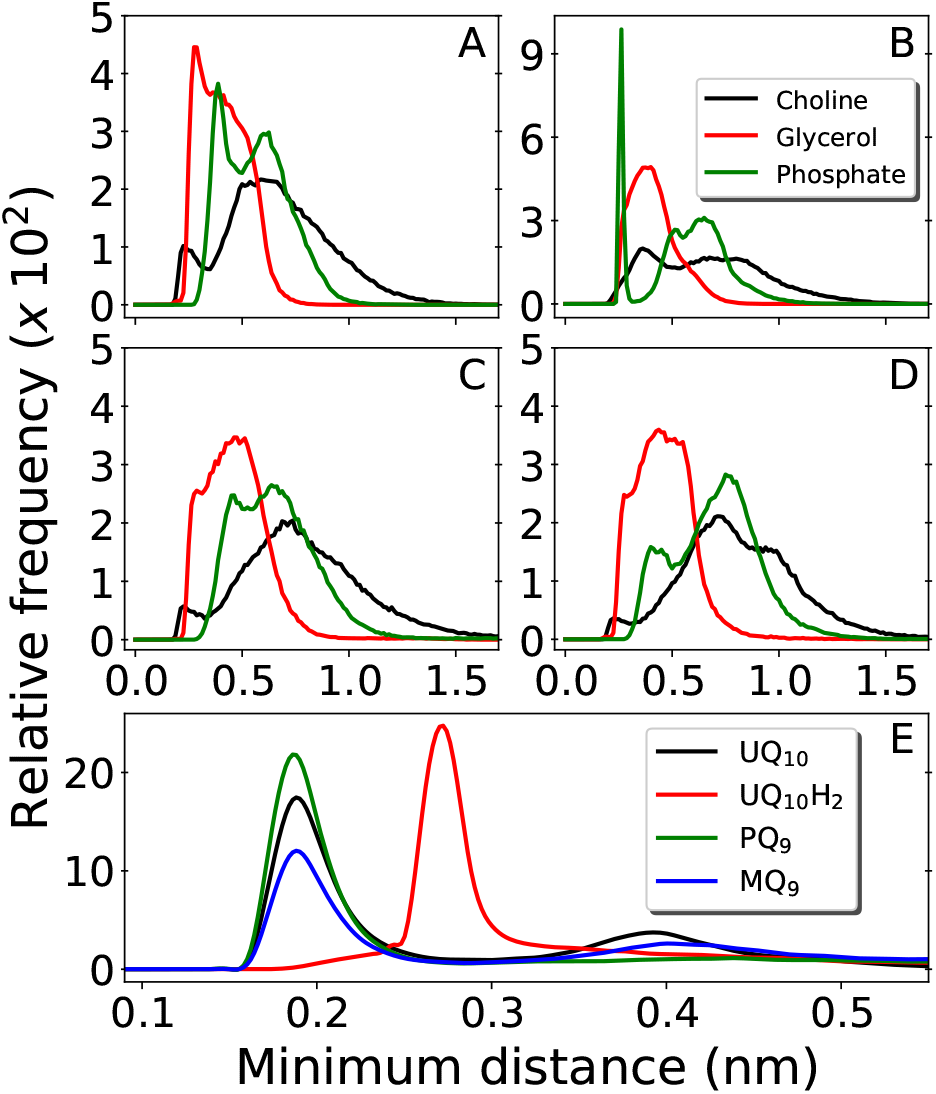
Distribution of minimum distances around the Q-head oxygen O1 of multicomponent bilayers with different Q composition. Contacts with DLPC choline group are shown in black, DLPC+DLPE glycerol in red and DLPC+DLPE phosphate in green. Panel A is UQ_10_, B is UQ_10_H_2_, C is PQ_9_ and D is MQ_9_. Panel E shows water contacts for UQ_10_ in black, UQ_10_H_2_ in red, PQ_9_ in green and MQ_9_ in blue. Data obtained from unrestrained MD simulations.

The H-bond donor UQ_10_H_2_ frequently makes an H-bond with the anionic phosphate group in DLPC+DLPE (peak at 0.26 nm at Fig. 3B) or in cardiolipins (not shown). H-bond acceptors (UQ_10_, PQ_9_ and MQ_9_) do not show frequent contacts with phosphates, neither interact closely with the positive choline groups in DLPC (Fig. 3A, C and D). Almost no H-bonds are observed between any of the Q molecules and either the ethylamine group in DLPE or the bridge glycerol in cardiolipins (not shown). Instead, the Q-head is located near acyl glycerol groups (Fig. 2A). This is very similar to the equilibrium location and contacts performed by Q in a POPC bilayer.^26^

Results for MQ_9_ suggest the two rings in the naphtoquinone head have similar average location across the membrane normal (Fig. 2D), but are probably orthogonal to the bilayer plane with O1 less solvated (pointing towards the membrane) and the opposite O4 more often in contact with water (pointing to the water phase). For PQ9, the Q-tail density has the lowest peak and spreads more evenly than other Q with long tails. Thus, PQ_9_ may have more conformational flexibility which explains the lack of secondary solvation peak (Fig. 3E), without disturbing the average Q-head location.

In comparison to our previous simulations of Q in POPC membranes,^26^ both density profiles and contacts performed by the Q-head do not change significantly in this multicomponent membrane. The Q-head preferentially resides at the bilayer interface, near the membrane normal occupied by either POPC or DLPC+DLPE glycerol groups (Fig. 2A). No specific contacts have been found between Q and polar lipid heads, except for H-bonds between ubiquinol and phosphate groups which are present in any phospholipid. The equilibrium distribution and location of Q on the membrane models studied so far result from the balance of hydrophobic interactions by the Q-tail and hydration of the Q-head, corresponding to a typical amphiphilic molecule.^55^

### Increased membrane permeability and penetration of water bound to the Q-head

Free energy profiles and analysis obtained from umbrella sampling simulations for UQ_2_ permeation across the membrane are presented in this section. An analogue with only two isoprenoid units was used because it is more efficient to sample computationally and reliable experimental partition coefficients are available for comparison.^56,57^

The calculated free energy profile in Fig. 4A shows three important aspects: The minimum across the membrane normal, which represents the equilibrium position of the Q-head, is located at 1.75 nm as expected from the Q-head density peak in Fig. 2A and close to the minimum observed for UQ_2_ permeation in the POPC bilayer (1.65 nm).^26^ The insertion or binding free energy for UQ_2_ in the multi-component membrane is −22±2 kJ/mol, in excellent agreement with experiment (Table 1) and with the binding free energy in POPC (−21 ±1 kJ/mol).^26^ However, the free energy barrier for UQ_2_ permeation in the multi-component membrane, given by the free energy difference between the minimum (at 1.75 nm) and the maximum near the center of the bilayer (0 nm), is only 19±1 kJ/mol, much lower than the barrier computed for permeation in POPC (47±2 kJ/mol).^26^

**Figure 4:**
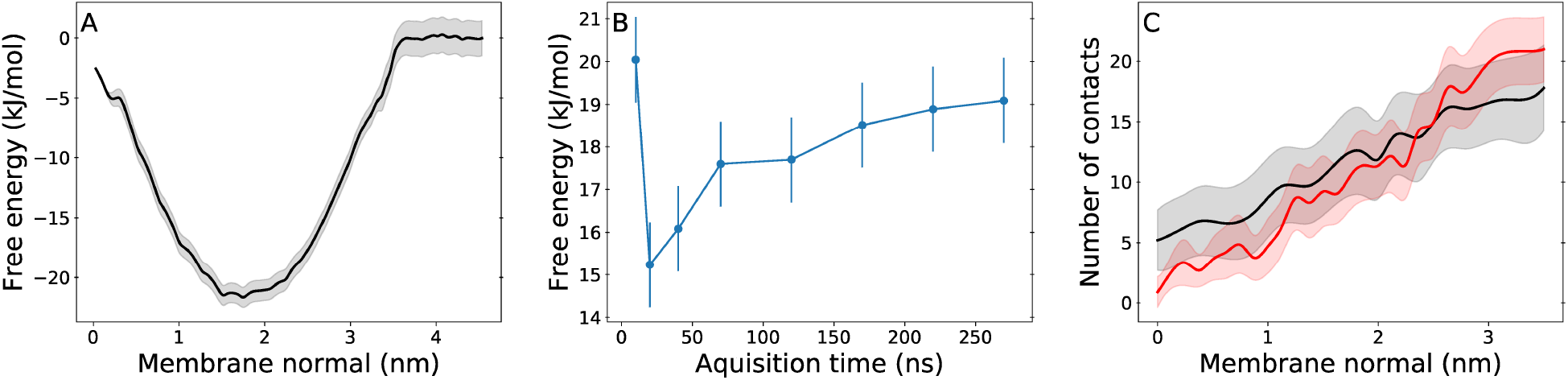
Free energy and hydration profiles for UQ_2_ permeation. Panel A shows the free energy profile with statistical uncertainty (gray shadow) for UQ_2_ insertion across the membrane normal. B shows the convergence of the permeation barrier with increasing acquisition time and fixed 30 ns of initial equilibration discarded from analysis. C shows the average number of water contacts with the Q-head for permeation in the multi-component membrane (black) and in the pure POPC membrane (red),^26^ with standard deviations shown in shadow. Data obtained from umbrella sampling simulations and symmetrized between the two leaflets.

**Table 1:** Experimental and simulated binding free energies (in kJ mol^−1^) between ubiquinone analogues and multi-component membranes. Experimental free energies were obtained from partition coefficients assuming a temperature of 310K. Values from Róg *et al*.^38^ were estimated from their Fig. 3.

| molecule | Experimental | Róg<sup>38</sup> | Here |
| --- | --- | --- | --- |
| UQ<sub>1</sub> | -17,<sup>57</sup> -20<sup>56</sup> | -34±4 |  |
| UQ<sub>2</sub> | -21,<sup>56</sup> -24<sup>57</sup> |  | -22±2 |

Fig. 4B shows the permeation barrier converges to within one statistical uncertainty in 200 ns of simulation (170 ns of data acquisition) per US window. In total, 50% more data (300 ns) were collected in each window, supporting the quality of the barrier computed here. Experimental estimates of rates for Q permeation or “flip-flop” by NMR linewidth^25^ and redox titration of substrates trapped inside vesicles^24^ only suggested rate upper bounds and for lipids with fully saturated acyl chains. To our knowledge, there are not measurements of Q transversal diffusion for bilayers composed of unsaturated acyl chains to compare with the barrier determined here.

Thus, Q permeation is observed more frequently in the multi-component membrane than in POPC. Although we have not computed profiles for permeation of the other Q studied here, spontaneous flip-flops were observed for UQ_2_H_2_, PQ_9_ and MQ_9_ at least once during the 300 ns of unrestrained MD simulations presented in the previous section. Given that UQ_*n*_ did not show spontaneous flip-flops, we can speculate that the other Q analogues will show smaller permeation barriers.

Then, what is different in this multicomponent membrane that allows permeation more frequently? The mean area per lipid heavy atom is 0.0126 nm^2^/atom in the multi-component membrane versus 0.0119 nm^2^/atom in POPC,^26^ suggesting the former membrane is less packed. The two unsaturations (positions 9 and 12) in the linoleoyl acyl chain composing all lipids in the multicomponent membrane result in a less ordered lipid phase, as shown by lower methylene order parameters computed for linoleoyl chain in comparison to palmitoyl and oleoyl in POPC (Fig. 5B). Lower lipid packing and ordering result in a more porous membrane in which molecules can penetrate with a cheaper cavitation energy cost.^22,58^

**Figure 5:**
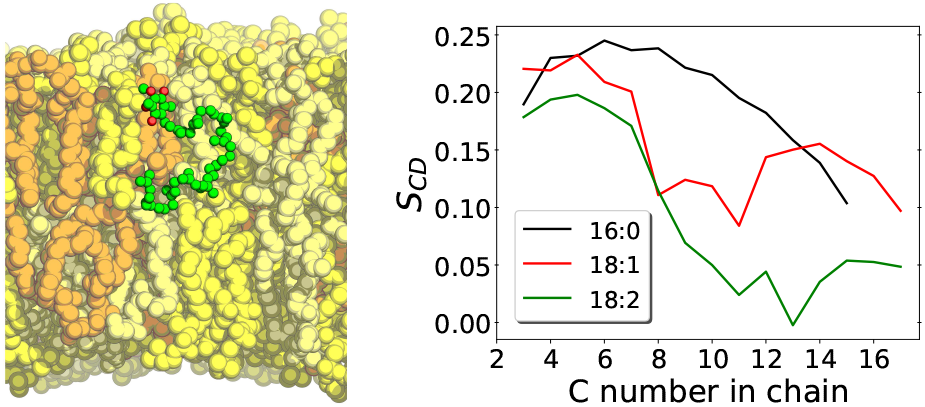
A) Left panel shows a structural snapshot of the multi-component membrane with UQ_10_ in its equilibrium position. LCL is shown in orange, DLPC in yellow, DLPE in beige and UQ_10_ in green (carbons) and red (oxygens). B) Right panel shows carbon-hydrogen bond vector order parameters (*S_CD_*) calculated with unrestrained MD for acyl chains POPC palmitoyl in black and oleoyl in red,^26^ and for DLPC+DLPE linoleoyl acyl chains in green.

Accordingly, more water contacts and H-bonds with the Q-head are performed inside the lipid phase for UQ_2_ permeation in the multi-component membrane (normal < 1 nm in Fig. 4C), leading to less desolvation of the polar Q-head and a lower permeation barrier than in POPC. Water arrest inside the lipid phase during the permeation of polar solutes has been observed in other simulations.^33,59^ It should be noted that no water penetration is observed in equilibrium for both POPC or multi-component membranes. But in the rare event of Q permeation, it is easier for water bound to the Q-head to penetrate into a membrane composed by fatty acids with a higher fraction of unsaturations.

A comparison with the simulations performed by Róg *et al*. shows large and important qualitative differences, specially for ubiquinone which was reported to make no contacts with solvent or lipid head groups and localize in the center of the bilayer.^38^ It seems unlike that the polar head group in UQ_*n*_, with four oxygen H-bond acceptors, will be stable in the middle of the lipid phase. Their calculated binding free energy for UQ_1_ is in large disagreement with experimental partition coefficients (Table 1). For ubiquinol, Q-head location and contacts are similar to ours, but strangely depend on the number of isoprenoid units. The lack of symmetry between the two leaflets in the density profiles shown and the irregular dependence of calculated densities and contacts with the number of isoprenoid units question whether appropriate equilibration and sampling were reached in their work.^38^ Exactly the same lipid composition and similar simulation procedures were employed here. Thus, most of the qualitative differences found should be due to force-fields. The OPLS derived force-field used by Róg *et al*. has been shown to artificially overbind Na+ cations^60^ and describe lipid order parameters in disagreement with NMR measurements.^61^ On the other hand, the CHARMM36 force-field used here has consistently one of the best performances describing these properties as well as several other membrane structural, thermodynamical and diffusional observables,^39,62,63^ including, of course, binding free energies and lateral diffusion constants for Q molecules.^26^

### Transversal diffusion of Q may limit turnover of electron transfer chains

Fig. 6 shows the equilibrium position of the Q-head in the multi-component membrane also matches the normal position (in modulus) of entrance sites for Q binding in respiratory complexes, as noted previously for the POPC bilayer.^26^ An initial encounter complex formed after membrane diffusion of Q and collision into a respiratory complex will frequently have the appropriate geometry in the membrane normal. Fig. 6 also shows membrane permeation has to occur for quinone/quinol recycling as their sites of reduction/oxidation are placed at opposite sides.

**Figure 6:**
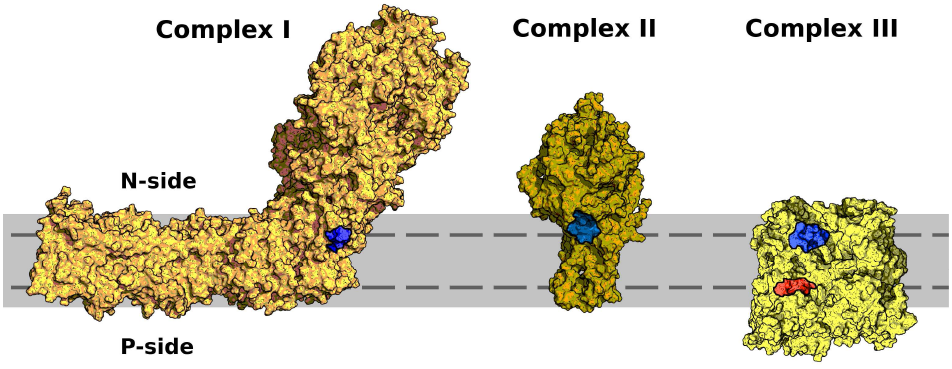
Entrance sites for Q binding in respiratory complexes I-III. Sites for quinone reduction shown in blue and for quinol oxidation in red. Bilayer is represented in solid gray and equilibrium or preferential position of the Q-head across the membrane normal (±1.75 nm) shown with a dashed line. Structures were taken from PDB ID 4HEA ^64^ for complex I, 2H88^65^ for complex II and 2QJP^66^ for complex III.

Bilayers with a low fraction of unsaturated chains, such as POPC,^26^ may slow down transversal diffusion by up to five orders of magnitude, in comparison to the bilayer with 2 unsaturations per chain studied here. This steep relation is due to the activated nature of the permeation process.^22,23^ A low fraction of unsaturation (<20%) may slow Q permeation up to a point it becomes rate-limiting for ETC or even blocks cellular growth.^13^ Yet another reason for the high content of cardiolipins, composed mostly by unsaturated acyl chains, in membranes involved in biological energy transduction^67^ might be the increment of Q permeability gained by lipid unsaturation.

## Conclusions

Molecular dynamics simulations for ubiquinone, ubiquinol, plastoquinone and menaquinone and free energy calculations for permeation of UQ_2_ in a multi-component bilayer mimicking the composition of bioenergetic membranes were presented.

Densities of Q-head and Q-tail (of similar length) across the membrane normal change little between each Q studied. The Q-head preferentially localizes near acyl glycerol groups and no specific contacts between the Q-head and polar lipid heads were observed, except for H-bonds formed between ubiquinol and phosphate groups. This is equivalent to the localization and contacts found for Q in a POPC bilayer. The binding free energy and equilibrium position estimated for UQ_2_ partition into the multi-component bilayer are also in agreement to what was found in POPC.^26^

We may conclude the equilibrium distribution of Q across model bilayers recently studied by molecular simulation^26,38^ do not change significantly with lipid composition, even when containing the anionic cardiolipin. The equilibrium structure and thermodynamics of embedded Q are determined by Q-head hydration and the hydrophobic effect of the isoprenoid tail. This should not be entirely unexpected given that similar partition coefficients have been measured for ubiquinone (with the same number of isoprenoid units) in biological membranes with varied composition.^56,57^

However, permeation of Q across the membrane changes considerably with lipid composition, specially the fraction of acyl chain unsaturation. Lipids with two unsaturations per acyl chain are less packed and less ordered, giving a more porous bilayer. Thus, the polar Q-head may carry water molecules during a flip-flop event with a lower free energy barrier, resulting in much faster transversal diffusion across the multi-component membrane studied here.

Membrane permeability modulated by the fraction of acyl chain unsaturation is important for efficient quinone/quinol recycling in opposite membrane sides and for translocation of protons by Q redox loops. Permeability may even limit turnover rates for biological ETC if the fraction of lipid unsaturation is too low. Results in the multi-component bilayer also confirm our previous observation that the equilibrium position of the Q-head across the membrane normal is similar to the position of protein sites for entrance of Q into respiratory complexes.^26^ Finally, it should be mentioned that lipid composition may also influence the structure of bioenergetic membranes^68^ and of respiratory complexes^69^ and their reactivity.^70,71^

## Acknowledgement

Funding from FAPESP (project 16/24096-5) and computational resources from the SDumont cluster in the National Laboratory for Scientific Computing (LNCC/MCTI) are gratefully acknowledged.

## Footnotes

† Abbreviations: Q is used in general for quinone or quinol analogues with any oxidation state and number of isoprenoid units. Q-head is the aromatic ring and bound groups except for the Q-tail, which is the full isoprenoid chain. UQ_*n*_ is oxidized ubiquinone, PQ_*n*_ is plas-toquinone and MQ_*n*_ is menaquinone, where the *n* subscript is the number of bound isoprenoid units. UQ_*n*_H_2_ is reduced ubiquinol. POPC is 16:0,18:1 phosphatidylcholine, DLPC is di-18:2 phosphatidylcholine, DLPE is di-18:2 phosphatidylethanolamine, LCL is tetra-18:2 cardiolipin, ETC is electron transfer chain, MD is molecular dynamics, US is umbrella sampling and COM is center-of-mass.

