## Supporting Information for "Effects of lipid composition on membrane distribution and permeability of natural quinones"

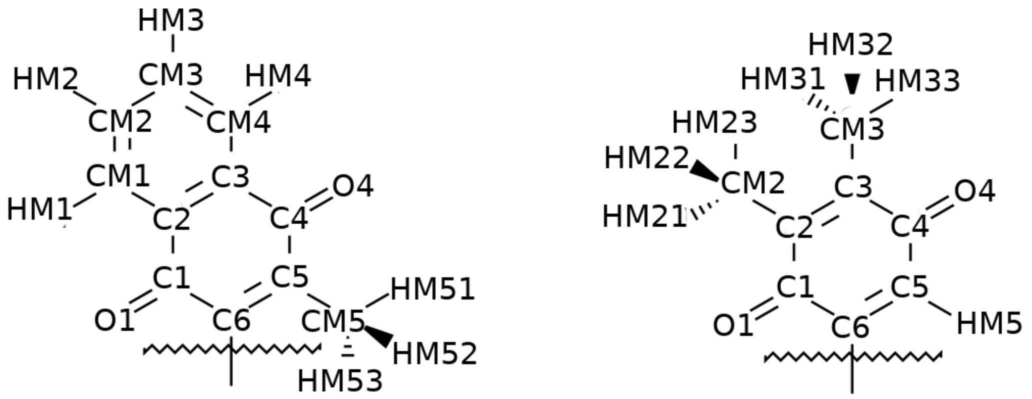


Figure S1: Atom names for PQ and MQ quinone heads.

Table S1: Atomic types and partial charges for PQ and MQ quinone heads. See figure S1 for atom names.

| **PQ head** | | |
|---|---|---|
| Atom name | Atom Type | Partial Charge |
| C5 | CA | -0.0575 |
| C6 | CA | -0.0575 |
| C4 | CA | 0.57 |
| O4 | O | -0.57 |
| C3 | CA | -0.115 |
| HM5 | HP | 0.115 |
| C2 | CA | -0.115 |
| C1 | CA | 0.57 |
| O1 | O | -0.57 |
| CM3 | CT3 | -0.155 |
| HM31 | HA3 | 0.09 |
| HM32 | HA3 | 0.09 |
| HM33 | HA3 | 0.09 |
| CM2 | CT3 | -0.155 |
| HM21 | HA3 | 0.09 |
| HM22 | HA3 | 0.09 |
| HM23 | HA3 | 0.09 |
| **MQ head** | | |
| Atom name | Atom Type | Partial Charge |
| C5 | CA | 0.09 |
| C6 | CA | 0.09745 |
| C4 | CA | 0.57 |
| O4 | O | -0.57 |
| C3 | CA | 0.00 |
| CM4 | CA | -0.115 |
| C2 | CA | 0.00 |
| CM1 | CA | -0.115 |
| C1 | CA | 0.57 |
| O1 | O | -0.57 |
| CM5 | CQ31 | -0.45745 |
| HM51 | HA3 | 0.09 |
| HM52 | HA3 | 0.09 |
| HM53 | HA3 | 0.09 |
| CM3 | CA | -0.115 |
| HM3 | HP | 0.115 |
| HM4 | HP | 0.115 |
| CM2 | CA | -0.115 |
| HM2 | HP | 0.115 |
| HM1 | HP | 0.115 |
